# Antisense oligonucleotides targeting the miR-29b binding site in the *GRN* mRNA increase progranulin translation

**DOI:** 10.1101/2022.01.12.476053

**Authors:** Geetika Aggarwal, Subhashis Banerjee, Spencer A. Jones, Yousri Benchaar, Jasmine Bélanger, Myriam Sévigny, Denise M. Smith, Michael L. Niehoff, Monica Pavlack, Ian Mitchelle S. de Vera, Terri L. Petkau, Blair R. Leavitt, Karen Ling, Paymaan Jafar-Nejad, Frank Rigo, John E. Morley, Susan A. Farr, Paul A. Dutchak, Chantelle F. Sephton, Andrew D. Nguyen

## Abstract

Heterozygous *GRN* (progranulin) mutations cause frontotemporal dementia (FTD) due to haploinsufficiency, and increasing progranulin levels is a major therapeutic goal. Several microRNAs, including miR-29b, negatively regulate progranulin protein levels. Antisense oligonucleotides (ASOs) are emerging as a promising therapeutic modality for neurological diseases, but strategies for increasing target protein levels are limited. Here, we tested the efficacy of ASOs as enhancers of progranulin expression by sterically blocking the miR-29b binding site in the 3’ UTR of the human *GRN* mRNA. We found 16 ASOs that increase progranulin protein in a dose-dependent manner in neuroglioma cells. A subset of these ASOs also increased progranulin protein in iPSC-derived neurons and in a humanized *GRN* mouse model. In FRET-based assays, the ASOs effectively competed miR-29b from binding to the *GRN* 3’ UTR RNA. The ASOs increased levels of newly synthesized progranulin protein by increasing its translation, as revealed by ribosomal profiling. Together, our results demonstrate that ASOs can be used to effectively increase target protein levels by partially blocking miR binding sites. This ASO strategy may be therapeutically feasible for progranulin-deficient FTD as well as other conditions of haploinsufficiency.

## Introduction

Progranulin is a lysosomal and secreted protein with pleiotropic effects, including promoting neuronal survival, neurite outgrowth, wound healing, tumor cell growth, and modulating inflammation (1,2). In humans, heterozygous *GRN* mutations cause frontotemporal dementia (FTD) due to progranulin haploinsufficiency (3,4) and reduced progranulin levels are a risk factor for Alzheimer’s disease (5-11). Therefore, increasing progranulin levels is a therapeutic goal for these forms of dementia (12,13). Gene therapy studies in mice provide proof of concept that restoring progranulin in heterozygous *Grn* mice improves FTD-associated neuropathology and behavioral deficits (14). Current therapeutic efforts are focused on small molecules that increase progranulin expression (15-18), gene therapies (14,19), monoclonal antibodies that modulate progranulin trafficking (20), and protein replacement (21). However, there are currently no approved therapies for progranulin-deficient FTD.

Antisense oligonucleotides (ASOs), short synthetic oligonucleotides used to modulate target RNAs, are emerging as a promising therapeutic modality for neurological diseases (22,23). The most commonly used ASO strategies involve RNase H1-mediated degradation of the target mRNA and modulation of splicing of the target mRNA (22). ASO-based strategies for increasing target protein levels are still relatively limited. Several strategies have been reported, including targeting upstream ORFs (uORFs) and regulatory elements in the 5’ UTR (24,25). However, these approaches are limited by the fact that not all genes possess uORFs and/or 5’ regulatory elements.

MicroRNAs (miRs) have been estimated to regulate more than 60% of all human proteins (26). These miRs typically bind to the 3’ UTR of target mRNAs and decrease protein levels through translational repression or mRNA decay (27,28). For progranulin, three miRs have been reported to negatively regulate progranulin protein levels: miR-29b, miR-107, and miR-659 (29-33). The binding sites of miR-29b and miR-659 have been mapped to the *GRN* 3’ UTR (29-32). Notably, the miR-659 binding site overlaps with the rs5848 SNP (31), which is a risk factor for FTD (31) and Alzheimer’s disease (9,34). *In silico* analyses predict that this SNP affects the binding affinity of miR-659 to the *GRN* mRNA, with the T allele having stronger binding and therefore greater translational repression. Consistent with this, individuals with the TT genotype (minor allele) at rs5848 have ∼30% decreased progranulin protein in brain tissue and in CSF (31,35), suggesting that miR-mediated modulation of progranulin levels contributes to risk of these forms of dementia. Thus, these miRs present as potential targets for increasing progranulin protein.

In the current study, we tested a therapeutic strategy for increasing progranulin protein levels by using ASOs to target the miR-29b binding site in the human *GRN* mRNA. We show that these ASOs effectively increase progranulin protein in cultured cells, in iPSC-derived neurons, and in mouse brains. Additionally, we determined the mechanism of action is that these ASOs displace miR-29b from its binding site and thereby de-repress translation, resulting in increased synthesis of progranulin protein.

## Results

### ASOs targeting the miR-29b binding site increase progranulin protein levels

Previous studies have reported that miR-29b inhibition increases progranulin protein levels in cells (29,30). We confirmed that treatment with a broad miR-29b inhibitor increases progranulin protein levels (Fig. 1A) in H4 human neuroglioma cells, which have moderate expression of miR-29b (Fig. S1A). Data from the Human miRNA Tissue Atlas (36) show that miR-29b is expressed in the human brain (Fig. S1B).

**Figure 1.**
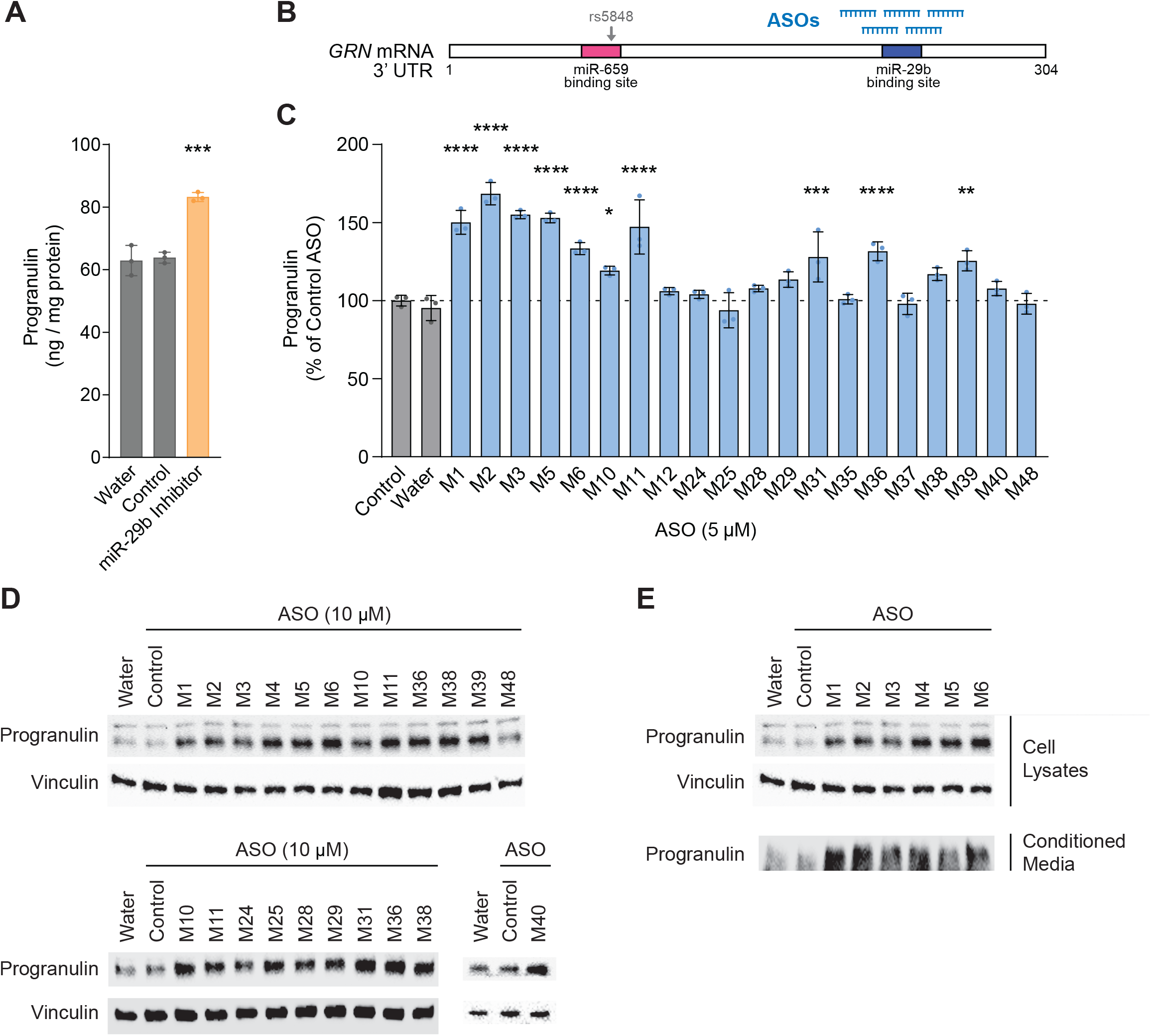
ASOs targeting the miR-29 binding site in the human *GRN* mRNA increase progranulin protein levels. *A*, ELISA for progranulin protein levels in H4 neuroglioma cells 24 h after transfection with 100 nM miR-29b inhibitor using Lipofectamine 3000. *B*, Schematic of the *GRN* 3’ UTR indicating miR-29b binding site and region targeted by ASOs. *C*, Validation of ASO hits from initial screening. H4 cells were treated with 5 μM ASO for 24 h, and progranulin levels were measured in cell lysates by ELISA. *D*, Western blot validation of candidate ASOs in H4 cells (10 µM, 24 h). *E*, ASOs increase both cellular and secreted progranulin levels. After treatment with 10 μM ASO for 24 h, progranulin levels in cell lysates and in the conditioned media were determined by western blot. Data are presented as means ± SD; * indicates p<0.05, ** indicates p<0.01, *** indicates p<0.001, **** indicates p<0.0001, as determined by one-way ANOVA with Dunnett post hoc test.

We next sought to test if ASOs targeting the miR-29b binding site in the *GRN* mRNA can increase progranulin protein levels. To this end, we designed a panel of 18-mer ASOs that span the entire miR-29b binding site with single nucleotide resolution (Fig. 1B). Our initial ELISA-based screening at 5 μM in H4 cells identified 11 ASOs that increased progranulin protein levels by >15% (Fig. 1C). We subsequently validated these 11 ASOs (M1, M2, M3, M5, M6, M10, M11, M31, M36, M38, M39) and identified 5 additional ASOs (M4, M25, M28, M29, M40) that increased progranulin protein levels at 10 μM, as determined by western blot (Fig. 1D). We retested a subset of these ASOs against a panel of negative control ASOs and found that the *GRN*-targeting ASOs (M3, M5, M10, M38) increased progranulin protein levels compared to all three negative control ASOs [two non-targeting control ASOs and a previously reported *MAPT*-targeting ASO (37)] (Fig. S2).

We noted that the efficacious ASOs largely clustered in two regions: 1) overlapping with the seed sequence (M25–M40), and 2) toward the 5’ end of the miR-29b binding site (M1–M6, M10–M11). Notably, ASOs M1–M5 do not overlap with the binding site. The predicted secondary structure of the *GRN* mRNA (Fig. S3) reveals that this region forms a large loop, which could increase accessibility for ASO binding. Moreover, this loop is in close proximity to the miR-29b binding site and may interact with the miR-29b binding site, which could in part explain the efficacy of ASOs M1–M5. Because progranulin is also secreted from cells, we tested if the ASOs similarly increase progranulin in the conditioned media. We found that ASOs that increased cellular progranulin levels also increased secreted progranulin levels (Fig. 1E), suggesting that the observed increases in cellular progranulin levels are not due to altered trafficking or secretion.

We then carried out time course and dose curve experiments for selected ASOs in H4 cells. The time course studies revealed that increased progranulin protein levels are detectable within 1–2 h of ASO treatment for ASOs M6 and M10, and by 8 h for ASO M36 (Fig. 2A). The slower kinetics of ASO M36 may be due its binding properties, as it has a lower melting temperature than ASOs M6 and M10. Dose curve experiments showed that the ASOs exhibit dose-dependent effects with EC_50_ values in the range of 1–8 μM (Fig. 2B). These ASOs exhibited no detectable toxicity in H4 cells, as determined by MTT assays (Fig. S4).

**Figure 2.**
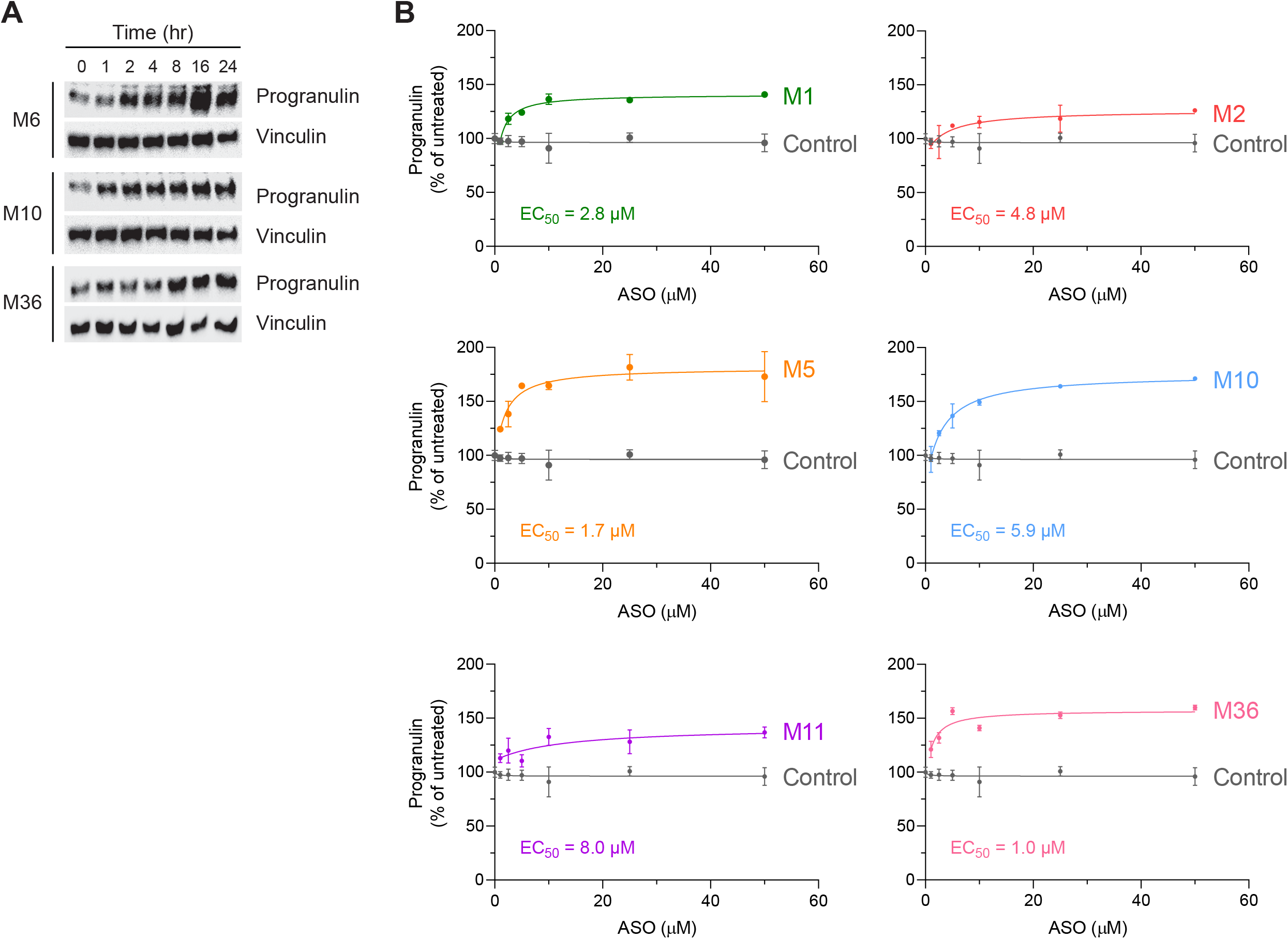
Time course and dose curve experiments for selected ASOs. *A*, ASOs increase progranulin protein levels within several hours. H4 cells were treated with 10 µM ASO for the indicated times, and progranulin levels in cell lysates were determined by western blot. *B*, Dose curve of ASO treatment in H4 cells (24 h). Progranulin levels were measured in cell lysates by ELISA. Data are presented as means ± SD, with EC_50_ values indicated.

We next tested selected ASOs in iPSC-derived cortical neurons and found that these ASOs similarly increased progranulin protein levels in a dose-dependent manner (Fig. 3A). To test the ASOs *in vivo*, we used a recently developed humanized *GRN* mouse model (38); intra-striatal injection of a miR-29b inhibitor was previously shown to increase progranulin protein levels in the brains of these mice (38). At three weeks following intracerebroventricular (ICV) administration of 500 μg ASO (39), we observed increased human progranulin protein by ELISA in the cortex of mice that received ASO M5 (Fig. 3B). Progranulin levels were increased by 63% and 61% in male and female mice, respectively, compared to mice of the same sex that received the scrambled control ASO. By western blot, we also observed increased human progranulin levels in the thalamus and hippocampus of male mice that received ASO M5 (Fig. 3C). Together, these results demonstrate that ASOs targeting the miR-29b binding site can increase human progranulin protein levels in cultured cells, neurons, and *in vivo*.

**Figure 3.**
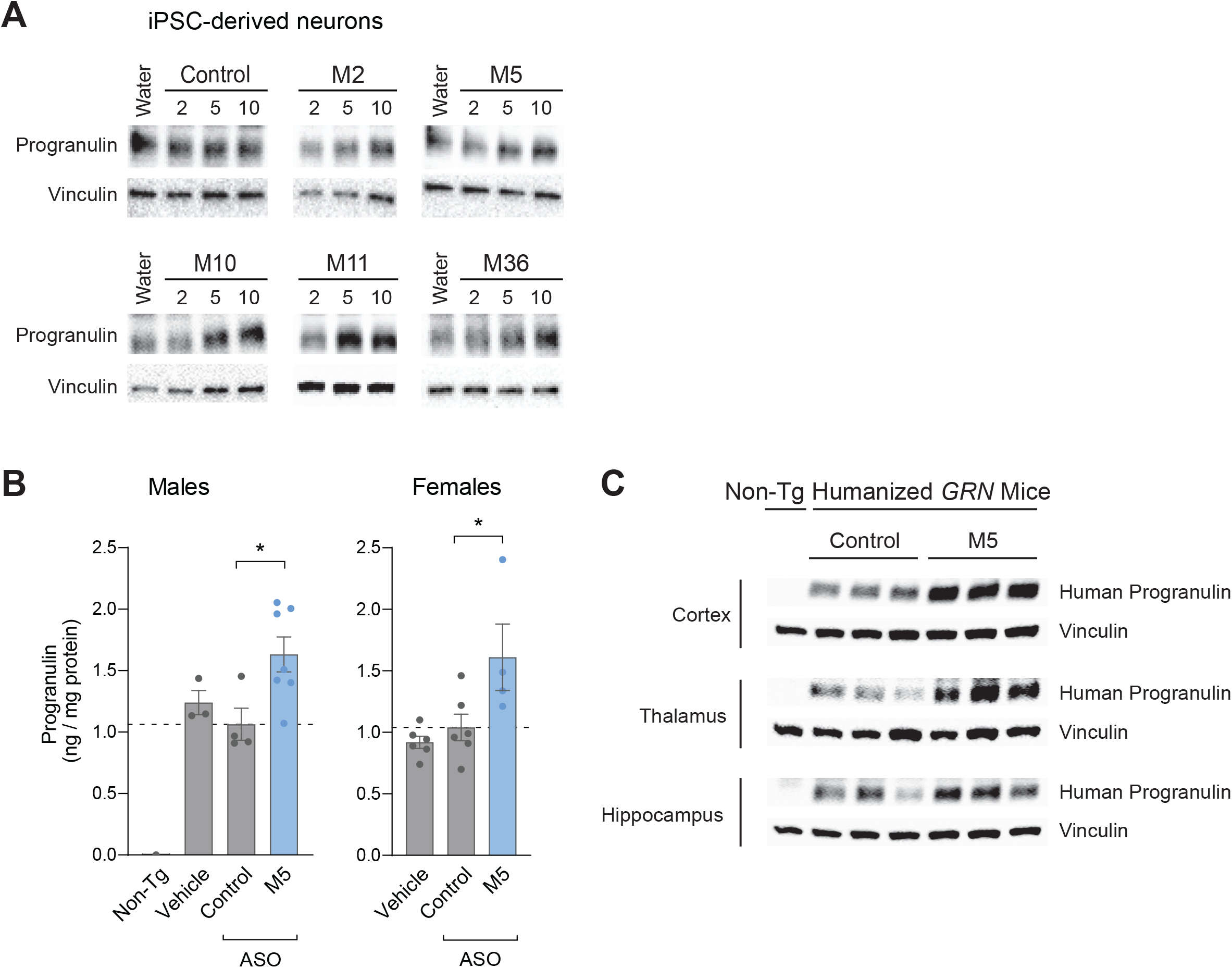
ASOs increase progranulin protein levels in iPSC-derived neurons and in mouse brains. *A*, Western blot of progranulin in iPSC-derived neurons following ASO treatment at the indicated concentrations (µM) for 3–4 days. *B*–*C*, Human progranulin levels in brains of humanized *GRN* mice at 3 weeks after ICV administration of 500 µg ASO. *B*, Human progranulin levels in cortex, as determined by ELISA. *C*, Western blot of human progranulin in cortex, thalamus, and hippocampus. Data are presented as means ± SEM; * indicates p<0.05, as determined by one-way ANOVA with Dunnett post hoc test. Non-Tg, non-transgenic.

### ASO effect requires miR-29b and involves competition of miR-29b from the GRN 3’ UTR

We next tested if the ASOs act through miR-29b by using a miR-29b inhibitor. This inhibitor binds to miR-29b and thereby prevents miR-29b binding to its target RNAs. The experiments of Fig. 4A demonstrate that ASOs M5 and M10 increased progranulin levels similarly to the miR-29b inhibitor and importantly that the miR-29b inhibitor abrogated any further effects of the ASOs on progranulin levels. These results strongly suggest that the ASOs block miR-29b’s effect on progranulin.

**Figure 4.**
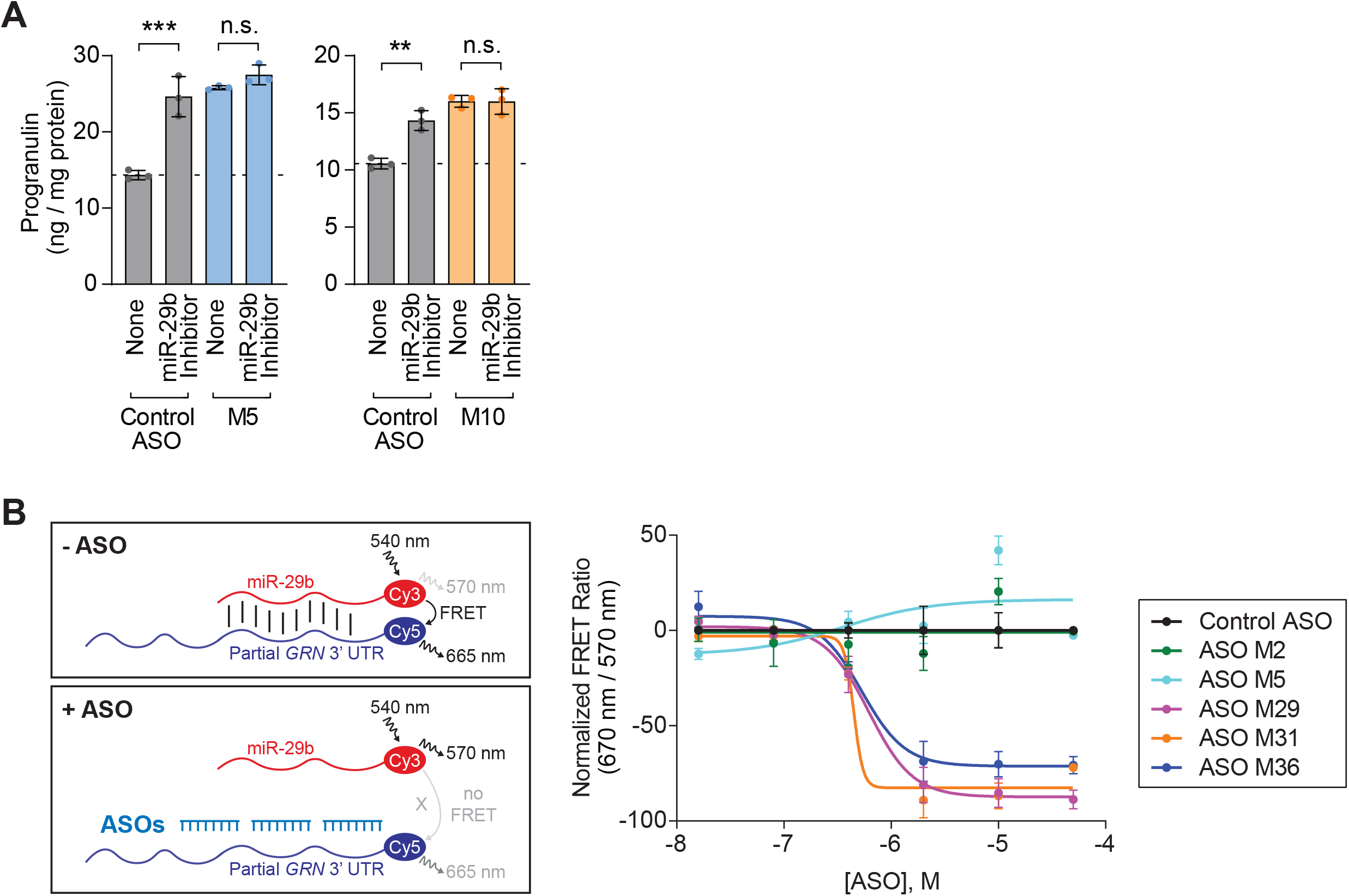
ASOs act through blocking miR-29b binding. *A*, ASOs require miR-29b to increase progranulin protein levels. ELISA for progranulin protein levels after co-treatment with miR-29b inhibitor (70 µM) and ASOs (20 µM) for 24 h. *B*, FRET assay demonstrating multiple ASOs can compete miR-29b from binding to a partial *GRN* 3’ UTR RNA. Data are presented as means ± SD; ** indicates p<0.01 and *** indicates p<0.001 as determined by two-way ANOVA with Tukey post hoc test. n.s., not significant.

To test if the ASOs can compete for binding of miR-29b, we used a FRET-based assay to monitor the *in vitro* interaction between miR-29b and a *GRN* 3’ UTR RNA that contains the miR-29b binding site. We found that several ASOs (M29, M31, and M36) were able to effectively compete off miR-29b binding (Fig. 4B), with IC_50_ values in the range of 0.4–0.8 µM. The scrambled control ASO and several ASOs which do not overlap with the binding site (i.e., M2 and M5) were unable to displace miR-29b; it is possible the partial *GRN* 3’ UTR RNA used in this assay does not completely recapitulate aspects of the full length *GRN* mRNA, such as the secondary structure (see Fig. S3). Together, these experiments indicate that the ASOs’ effects require miR-29b and involve competition for miR-29b binding to the *GRN* mRNA.

### ASOs increase translation of progranulin

Despite increasing progranulin protein levels, the ASOs did not increase *GRN* mRNA levels in H4 cells (Fig. 5A), suggesting that the ASOs increase the rate of progranulin translation. This would be consistent with the canonical effects of translational repression by many miRs (27,28). To formally test this, we performed ribosomal profiling experiments to assess the amount of ribosome-bound *GRN* mRNA, which reflects its rate of translation. We found that cells treated with ASOs M10 and M36 had marked enrichment of *GRN* mRNA in the heavy polyribosome fractions (11–16), compared to cells treated with a scrambled control ASO (Fig. 5B), indicating that these ASOs increase the rate of progranulin translation. This effect was specific to *GRN* mRNA, as it was not observed with *FUS* mRNA (data not shown), which is linked to amyotrophic lateral sclerosis (ALS) and FTD.

**Figure 5.**
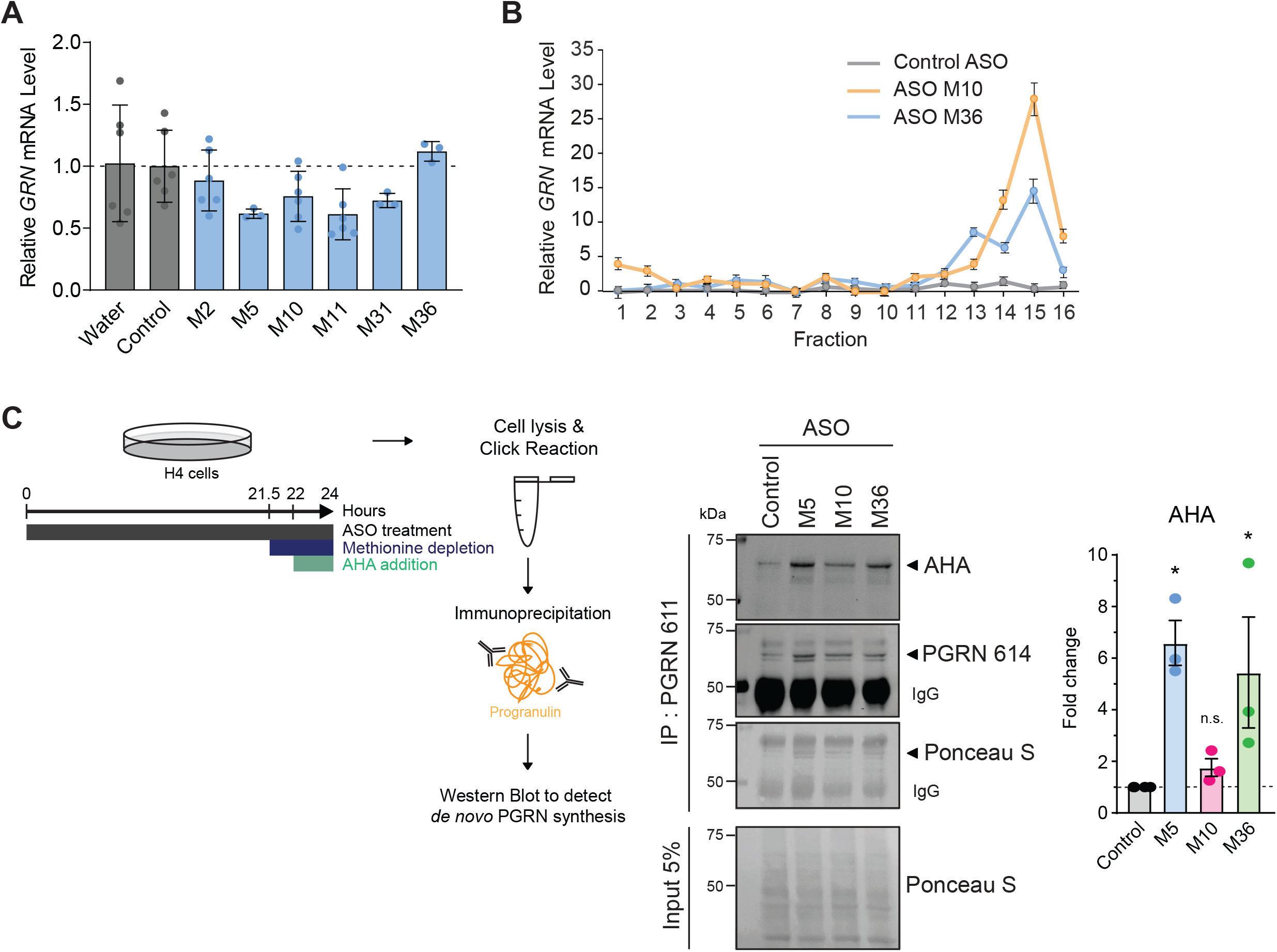
ASOs increase progranulin translation. *A, GRN* mRNA levels following ASO treatment (5 µM, 24 h), as determined by qPCR. *B, GRN* mRNA distribution in each of 16 sucrose gradient fractions from ASO-treated H4 cells (5 µM, 22 h) analyzed by qPCR. Fraction 1: hydrosoluble fraction; fractions 2, 3, and 4: 40S, 60S, and monosomes, respectively; fractions 5–10: light polyribosomes; and fractions 11–16: heavy polyribosomes. Data are plotted as a fraction of total mRNA on the gradient. *C*, Western blot showing immunoprecipated, newly synthesized, AHA/biotin-labeled progranulin protein following ASO treatment (10 µM, 24 h). Data are presented as means ± SD; * indicates p<0.05 and ** indicates p<0.01, as determined by one-way ANOVA with Dunnett post hoc test. n.s., not significant. PGRN, progranulin.

To determine if the ASOs increase the amount of newly synthesized progranulin protein, we used the methionine analog azidohomoalanine (AHA) to label nascent proteins. Following immunoprecipitation of progranulin, western blots using fluorescently labeled-streptavidin confirmed that ASOs M5 and M36 increase synthesis of progranulin protein (Fig. 5C). Together, these studies establish the mechanism of action is that these ASOs displace miR-29b from its binding site and thereby de-repress translation, resulting in increased synthesis of progranulin protein.

## Discussion

In the current study, we show that ASOs targeting the miR-29b binding site can increase human progranulin protein levels in cultured cells, in iPSC-derived neurons, and in the brains of humanized *GRN* mice. In cells, the ASO effects are dose dependent and are detectable within several hours. Our mechanistic studies revealed that the ASOs act through miR-29b, compete for miR-29b binding to the 3’ UTR of the *GRN* mRNA, increase the rate of progranulin translation, and ultimately increase the amount of newly synthesized progranulin protein. To our knowledge, this is the first in-depth characterization of using ASOs to increase translation of a target protein by sterically blocking a miR binding site. Several previous studies used oligonucleotides to block miR binding sites (40-45), but to our knowledge this is the first report using this strategy to increase progranulin levels.

This ASO-based strategy has several notable advantages. The increased progranulin protein is endogenously produced from the intact *GRN* allele, and therefore is expected to have identical properties and function; this avoids unexpected complications from ectopic *GRN* expression and/or modified forms of recombinant progranulin used in other therapeutic strategies. Additionally, this strategy increases progranulin protein levels within a narrow physiological window and thus would avoid issues arising from high level overexpression of progranulin, such as possible tumorigenesis (46,47). Moreover, by targeting regions partially overlapping with the miR-29b binding site in the *GRN* mRNA increases the specificity of our approach by avoiding other miR-29b target genes due to sequence differences between genes in the regions flanking the binding site. Lastly, this ASO strategy targeting the miR-29b binding site is agnostic to the specific disease mutation and could be used in the context of any of the more than 70 FTD-associated *GRN* mutations that have been identified (48).

Several lines of evidence support that the increases in progranulin protein levels observed following treatment with the *GRN*-targeting ASOs are the result of on-target effects mediated by sterically blocking miR-29b binding to its binding site in the *GRN* 3’ UTR. First, multiple ASOs are able to physically compete for miR-29b binding to the *GRN* 3’ UTR RNA in a FRET-based assay (Fig. 4B). Second, the ASOs’ effects are dependent on miR-29b, as the ASOs do not increase progranulin protein levels when miR-29b is blocked (Fig. 4A). Furthermore, the ∼70% maximal increases in the progranulin levels observed with the *GRN*-targeting ASOs in H4 cells (ASOs M5 and M10 in Fig. 2B) are comparable to the ∼70% maximal increases observed with a broad miR-29b inhibitor (Fig. 4A). Third, in ribosomal profiling experiments, the ASOs specifically increase the rate of *GRN* translation (Fig. 5B); such an effect was not observed with *FUS* mRNA (data not shown), which is linked to amyotrophic lateral sclerosis (ALS) and FTD. Fourth, *in silico* analysis found no predicted off-target RNAs in the human genome with zero mismatches for any of the ASOs used in this study. Further analyses allowing for 1 bp mismatch revealed a total of seven potential off-target RNAs within exonic sequences; however, three are not expressed in the human brain based on the GTEx Portal (49), and none of the remaining four potential targets are shared by multiple ASOs. Lastly, this effect on progranulin protein levels was observed with multiple efficacious *GRN*-targeting ASOs (16 in total), when compared to three negative control ASOs (Fig. S2) and many non-efficacious *GRN*-targeting ASOs with identical chemistry (Fig. 1C–D), further strengthening the case that these are on-target effects.

It is unclear how ASOs M1–M5 increase progranulin levels, despite not overlapping with the binding site. Based on the predicted secondary structure of the *GRN* mRNA (Fig. S3), the region targeted by ASOs M1–M5 harbors a large loop, which could provide increased accessibility for ASO binding. Moreover, this loop is in close proximity to the miR-29b binding site and may interact with the miR-29b binding site, which could in part explain the efficacy of ASOs M1–M5. Our data suggests that, similar to ASOs that overlap with the binding site, ASO M5 increases progranulin translation (Fig. 5C) and acts in a miR-29b-dependent manner (Fig. 4A). In our *in vitro* FRET-based assay, ASOs M2 and M5 did not compete miR-29b from the *GRN* 3’ UTR; one possible reason is that the partial *GRN* 3’ UTR RNA used in this assay may not have the same secondary structure as the full length *GRN* mRNA.

Overall, ASOs are proving to be a promising therapeutic modality for CNS diseases, as ASOs are versatile modulators of endogenous RNAs, well tolerated in humans, and stable in the CNS with long duration of action (22,23). Future studies will focus on testing if these ASOs targeting the miR-29b binding site can delay or prevent FTD-associated neuropathology and behavioral deficits in mouse models. While the current study focuses on ASOs that target the miR-29b binding site, we have also tested a smaller panel of ASOs targeting the miR-659 binding site (31,32) in the *GRN* 3’ UTR (see Fig. 1B) and identified ASOs that similarly increase progranulin protein levels (Fig. S5A). Moreover, ASOs targeting the miR-659 and miR-29b binding sites have additive effects (Fig. S5B); this may provide an opportunity to further increase progranulin *in vivo*. Our results suggest that this ASO strategy may be broadly applicable for pharmacologically increasing levels of target proteins and could be particularly useful for development of ASO-based therapies for diseases of haploinsufficiency.

### Experimental procedures

#### ASOs

ASOs used in these studies were 18-mer ASOs, except the miR-29b inhibitor is a 23-mer ASO. ASOs used for in vitro and cell-based studies were dissolved in water and stored at - 20 °C. For *in vivo* studies, lyophilized ASOs were dissolved in sterile PBS without calcium or magnesium (Gibco, 14190-250) and sterilized by passing through a 0.2 µm filter. Off-target candidate genes with complementary RNA sequences were predicted using GGGenome with RefSeq human RNA release 205 (March 2021).

#### Cell culture

H4 human neuroglioma cells were obtained from ATCC (HTB-148) and cultured in DMEM (Dulbecco’s Modified Eagle Medium, high-glucose) (Gibco, 11995-073) supplemented with 10% fetal bovine serum (FBS) (Gibco, 26140-095), 10 U/ml penicillin, and 10 µg/ml of streptomycin. For typical ASO treatments, cells were plated in 6-well or 12-well plates, and then treated with ASOs as indicated on the following day. Human iPSCs harboring doxycycline-responsive Neurogenin-2 (NGN2) expression for differentiation into i3 cortical neurons were kindly provided by Michael Ward. Cells were grown and differentiated as described (50). After 2 weeks of differentiation, i3 neurons were treated with ASOs for 3–4 days. All ASO treatments were by free uptake (gymnotic delivery), and all cells were maintained at 37 °C and 5% CO_2_.

#### ELISA and immunoblot analysis

For progranulin measurements, cells were rinsed with PBS and then lysed in RIPA buffer containing protease inhibitors (Roche, cOmplete Mini EDTA-free Protease Inhibitor Cocktail). Cleared lysates were transferred to new tubes, and protein concentrations were determined using the Bio-Rad DC Protein Assay Kit II. For ELISA, progranulin concentrations were determined in duplicate using 10–15 µl of lysates per well (typically 8–20 µg of total protein per well) using a sandwich ELISA assay (R&D Systems, DPGRN0). For experiments analyzing secreted progranulin, conditioned media was collected and cleared at 10,000 x g for 10 min at 4° C. For western blot analysis, sample buffer was added to the lysates or conditioned media, and the samples were heated at 95 °C for 10 min. Equal amounts of protein lysates (10–20 µg) or equal volumes of conditioned media (10–20 µl) were separated on SDS–PAGE gels. Proteins were transferred to nitrocellulose membranes using the Bio-Rad Turbo-Blot transfer system. After blocking and antibody incubations, membranes were incubated with SuperSignal West Pico or Femto enhanced chemiluminescent HRP substrate (ThermoFisher) and visualized using a Chemi-Doc system (Bio-Rad). Primary antibodies used for immunoblot analysis include: an anti-human progranulin linker 5 polyclonal antibody #614 that recognizes an epitope between amino acids 497–515 (51), an anti-human vinculin monoclonal antibody (Cell Signaling Technology, 13901), and anti-mouse progranulin polyclonal antibody (R&D Systems, AF2557), and an anti-α-tubulin monoclonal antibody (Sigma, T9026). The HRP-conjugated secondary antibodies used were goat anti-rabbit IgG (H+L) (Jackson Immuno Research Labs, 111-035-144), donkey anti-mouse IgG (H+L) (Jackson Immuno Research Labs, 715035150), and donkey anti-sheep IgG (H+L) (Jackson Immuno Research Labs, 713035147).

#### RNA analysis

Total RNA was isolated from cultured cells using the RNeasy Mini kit (Qiagen, 74106) with on-column DNase digestion (Qiagen, 79256). RNA was reverse-transcribed to obtain cDNA using the iScript cDNA synthesis kit (Bio-Rad, 1708891), and qPCR was performed using PowerUp SYBR Green Master Mix (ThermoFisher, A25777) with a Bio-Rad CFX384 Real-Time System. Primers sequences for human genes were as follows: CYCLO-F, GGAGATGGCACAGGAGGAAA; CYCLO-R, CCGTAGTGCTTCAGTTTGAAGTTCT; human GRN-F, AGGAGAACGCTACCACGGA; GRN-R, GGCAGCAGGTATAGCCATCTG; Results for qPCR were normalized to the housekeeping gene *CYCLO*, and evaluated by the comparative C_T_ method. For RNA-seq analysis of miR levels, the small RNA fraction was isolated from triplicate 100-mm dishes of H4 cells using the mirVana miRNA Isolation Kit (Invitrogen, AM1561). Libraries were constructed using the Ion Total RNA-Seq Kit v2 (Life Technologies, 4475936) and sequenced on an Ion Torrent Proton at the Saint Louis University Genomics Core Facility. Reads were aligned to the human genome hg19 assembly, and data are presented as total normalized nucleotide coverage per miRNA. Data on miR expression levels in human tissues was from the Human miRNA Tissue Atlas (36). The predicted secondary structure of the human *GRN* mRNA was generated using the mfold program (52).

#### Polyribosome profiling

H4 cells were cultured in 100-mm dishes and treated with 5 µM ASOs for 22 h. Five minutes prior to collecting cells, cycloheximide (CHX) was added to the cell culture media at a final concentration of 0.1 mg/ml. Cells were rinsed with ice-cold PBS, and then lysed in Polyribosome Lysis Buffer containing 20 mM Tris-HCl pH 7.4, 100 mM KCl, 5 mM MgCl_2_, 1% NP-40, 0.1 mg/ml CHX, 1 mM DTT, 0.02 U/ml SUPERase In RNase Inhibitor (Invitorgen, AM2694), protease inhibitors (Roche, cOmplete Mini EDTA-free Protease Inhibitor Cocktail), and phosphatase inhibitors (Roche, PhosSTOP) in nuclease-free water (Invitrogen, AM9937). Lysates were centrifuged (10,000 x g, 10 min, 4 °C), and the supernatant fractions were collected. Protein determination was performed using the Bio-Rad DC Protein Assay Kit II, and 1.35 mg per sample was loaded on a 15% to 45% sucrose density gradient (w/w sucrose, 20 mM Tris-HCl pH 7.4, 100 mM KCl, 5 mM MgCl_2_, Ultrapure water). Samples were centrifuged for 2 h at 210,200 x g at 4 °C in a SW41 Ti rotor. Polyribosome profiling was performed using a BR-188 density gradient fractionation system (Brandel) (sensitivity setting of 1, baseline setting of 20, and flow rate of 1.5 ml/min) with upward displacement and continuous monitoring at 254 nm using a UA-6 detector. Polyribosome (13 fractions) and monosome (3 fractions) fractions were collected in a volume of 600 µl, and the fractions were separated in two tubes (300 µl per tube). For subsequent RNA extraction, 600 µl Trizol Reagent (Invitrogen, 15-596-018) was added to each polyribosome fraction (300 µl), and samples were stored at −80 °C. RNA was extracted by adding 200 µl chloroform (Fisher, C298) to each fraction and centrifuging at 18,213 x g for 15 min. The aqueous phase was transferred to a tube containing 500 µl of isopropanol (Fisher, A416P-4) and 5 µg of glycogen (Sigma, G1767-1VL). Then samples were pelleted at 18,213 x g for 15 min, dried, washed with 70% ethanol, and centrifuged at 18,213 x g for 15 min. The pellet was dried and resuspended in 20 µl of Ultrapure water. Samples were digested with DNase (Roche) for 40 min at 37 °C. cDNA was generated using 2 µg of RNA for each fraction with a High-Capacity cDNA Reverse Transcription Kit (ThermoFisher, 4368814). qPCR was performed using PowerUp SYBR Green Master Mix (ThermoFisher, A25776) using a QuantStudio 5 Real-Time PCR System. Primers sequences were as follows: human Actin-F, TCCGTGTGGATCGGCGGCTCCA; human Actin-R, CTGCTTGCTGATCCACATCTG; human GRN-F, AGGAGAACGCTACCACGGA; and human GRN-R, GGCAGCAGGTATAGCCATCTG. Relative RNA levels were calculated using a C_T_ mean value normalized with an internal control (Actin). The delta C_T_ value is then normalized to the delta C_T_ value of the input (taken before the polyribosome profiling) from the corresponding condition.

#### AHA labeling

H4 cells were cultured in DMEM high glucose (Gibco, 11965-092) supplemented with 10% FBS (Gibco, 10010-023). Cells were cultured in 60-mm plate until about 80% confluence and treated with 10 µM ASOs for a total of 24 h. For optimal incorporation of the amino acid azidohomoalanine (AHA), cells were methionine-deprived for 2.5 h in methionine-free medium (DMEM without methionine, glutamine, and cysteine (Gibco, 21013024), supplemented with 0.5% FBS, 1X GlutaMAX (Gibco, 35050061), and 63 μg/ml L-Cysteine) containing 10 µM ASO. AHA was then added to the cell culture medium at a final concentration of 50 µM for the last 2 h of treatment to label de novo proteins. Cells were washed once with ice-cold PBS containing 0.1 mg/ml CHX, and then lysed in Triton lysis buffer containing 1% Triton X-100, 25 mM sodium pyrophosphate, 150 mM NaCl, 50 mM NaF, 5 mM EDTA, 5mM EGTA, 0.5% sodium deoxycholate, 20 mM HEPES pH 7.4, 10 mM β-glycerophosphate, 1 mM Na_3_VO_4_, 0.1 mg/ml CHX, and protease inhibitors (Roche, cOmplete Mini EDTA-free Protease Inhibitor Cocktail). Lysates were centrifuged (16,260 x g, 10 min, 4 °C), and the supernatant fractions were collected. Click reactions were performed using Click-iT Protein Reaction Buffer Kit (Invitrogen, C10276) according to the manufacturer’s protocol in order to label the AHA-incorporated proteins with biotin alkyne (ThermoFisher, B10185). The Click reaction samples were incubated on a rotator at 4 °C for 1 h, and diluted in lysis buffer at a 1:2 ratio before being subjected to anti-progranulin immunoprecipitation. For immunoprecipitation, the lysates were incubated with rotation with 8.4 μg of anti-human progranulin linker 4 polyclonal antibody #611 (which recognizes an epitope between amino acids 422–440) (51) overnight at 4 °C, followed by a 2 h incubation with Protein A Agarose beads (Roche, 11719408001) at 4 °C. Beads were washed three times with the lysis buffer and the immunoprecipitates were eluted, denatured and boiled (5 min, 95 °C) in 2X Laemmli buffer. For western blot analysis, the immunoprecipitated samples were resolved on 10% SDS–PAGE gels, transferred to nitrocellulose membranes, and stained with Ponceau. After blocking incubations, the membranes were incubated with IRDye 800CW Streptavidin (LICOR, 926-32230) to detect AHA-labelled proteins with the LICOR Odyssey imaging system. The membrane was reprobed with anti-human progranulin linker 5 polyclonal antibody #614 (1:3000) using Quick Western Kit-IRDye 680RD secondary antibody (LICOR, 926-69100).

#### MTT assay

MTT assays were used to assess the effects of ASOs on cell viability. H4 cells were plated in 96-well plates at a density of 5,000 cells per well. The following day, cell were treated with 10 µM ASO. After 21 h treatment, the media was removed and cells were incubated in 100 µl of 0.5 mg/ml thiazolyl blue tetrazolium bromide (MTT) (Sigma, M5655) solution prepared in fresh culture media for 4 h. After incubation, the MTT solution was removed and 100 µl of DMSO was added to dissolve the crystals formed. The absorbance was measured at 570 nm using a BioTek Synergy H1 plate reader. Experiments were performed with triplicate wells, and the percent viability was normalized to cells treated with vehicle only (water).

#### FRET-based assay for miR-29b binding

A FRET-based assay was developed to monitor the interaction between miR-29b (labeled with Cy3 at the 5’ end) and a partial *GRN* 3’ UTR RNA (nucleotides 189–246, labeled with Cy5 at the 3’ end). Reactions were set up in duplicate wells in black 384-well plates with 1 µM miR-29b and 1 µM *GRN* 3’ UTR in 20 µl of 1X SSC buffer, pH 7.0 containing 0.005% Tween-20. ASOs were added to the reactions in the same buffer at final concentrations between 0.64 nM–50 µM. Plates were read using a BioTek Synergy H1 plate reader with excitation at 540 nm and emission at 570 nm, and with excitation at 540 nm and emission at 665 nm. The ratio of fluorescence emissions at 665 nm / 570 nm was calculated for each well, and data were transformed with baseline correction.

#### Mouse studies

Animal procedures were approved by the Institutional Animal Care and Use Committee of Saint Louis University and followed NIH guidelines. Mice were housed in a pathogen-free barrier facility with a 12-h light/12-h dark cycle and provided food and water ad libitum. Humanized *GRN* mice (38) were on the C57BL/6J background and were genotyped either by PCR as described (38) or by real-time PCR (Transnetyx). For ICV ASO delivery, 500 µg ASO (in 5 µl PBS) was administered by intracerebroventricular (ICV) bolus injection into the right lateral ventricle of mice anesthetized with isoflurane, as previously described (53). After 3 weeks, mice were sacrificed and brain tissues were collected for protein and RNA analyses. For ELISA, progranulin levels were determined in duplicate using 150 µg of protein lysates per well using a sandwich ELISA assay (abcam, ab252364). Western blot and qPCR analyses were carried out as described above.

#### Statistical analyses

Data are presented as means ± SD or as means ± SEM, as indicated in the figure legends. Data were analyzed with GraphPad Prism software using the statistical tests described in the figure legends. P values < 0.05 were considered significant.

## Data availability

The RNA-seq data used in this study are available in the Gene Expression Omnibus (GEO; http://www.ncbi.nlm.nih.gov/geo/) under accession code GSE188498. All the other data supporting the findings of this study are available within the article and its supporting information files.

## Acknowledgements

We thank Michael Ward for providing iPSCs for differentiation into i3 neurons; the Genomics Core Facility at Saint Louis University for RNA-seq; Sophie Chang for technical assistance; and Laura Mitic, Robert Farese, Jr., Tobias Walther, Thi Nguyen, and members of the lab for helpful discussions.

## Figure Legends

**Figure S1. miR-29b expression levels in cells and tissues**. *A*, miR-29b is moderately expressed in human H4 neuroglioma cells. Ranked profile of miR expression levels, as determined by RNA-seq. miR-29b is indicated in red. *B*, Expression levels of miRs known to regulate progranulin levels in the human brain. Data from the Human miRNA Tissue Atlas (36).

**Figure S2. *GRN*-targeting ASOs increase progranulin protein levels compared to multiple negative control ASOs**. H4 cells were treated with 10 μM ASO for 24 h, and progranulin levels were determined by western blot.

**Figure S3. Predicted secondary structure of the human *GRN* mRNA shows a large accessible loop immediately upstream of the miR-29b binding site**. The mfold program was used to generate the predicted secondary structure. The miR-29b binding site is highlighted in yellow. The nucleotide numbering is based on NCBI reference sequence NM_002087.4, and the 3’ UTR includes nucleotides 1823-2126.

**Figure S4. ASOs are not toxic in cells**. H4 cells were treated with 10 μM ASO for 21 h, and then cell viability was assessed by MTT assay. Data are presented as means ± SD of three independent experiments.

**Figure S5. Several ASOs targeting the miR-659 binding site in the *GRN* 3’ UTR increase progranulin protein levels**. *A*, H4 cells were treated with 5 μM ASO for 24 h, and progranulin levels were measured in cell lysates by ELISA. *B*, H4 cells were treated with 80 μM total ASO (40 μM for each ASO plus 40 μM control ASO, or 80 μM control ASO) for 24 h, and progranulin levels were measured in cell lysates by ELISA. Data are presented as means ± SD; * indicates p<0.05 and ** indicates p<0.01, as determined by one-way ANOVA with Dunnett post hoc test.

