## Supplementary figures and images for "Antisense oligonucleotides targeting the miR-29b binding site in the *GRN* mRNA increase progranulin translation"

### Fig. S1

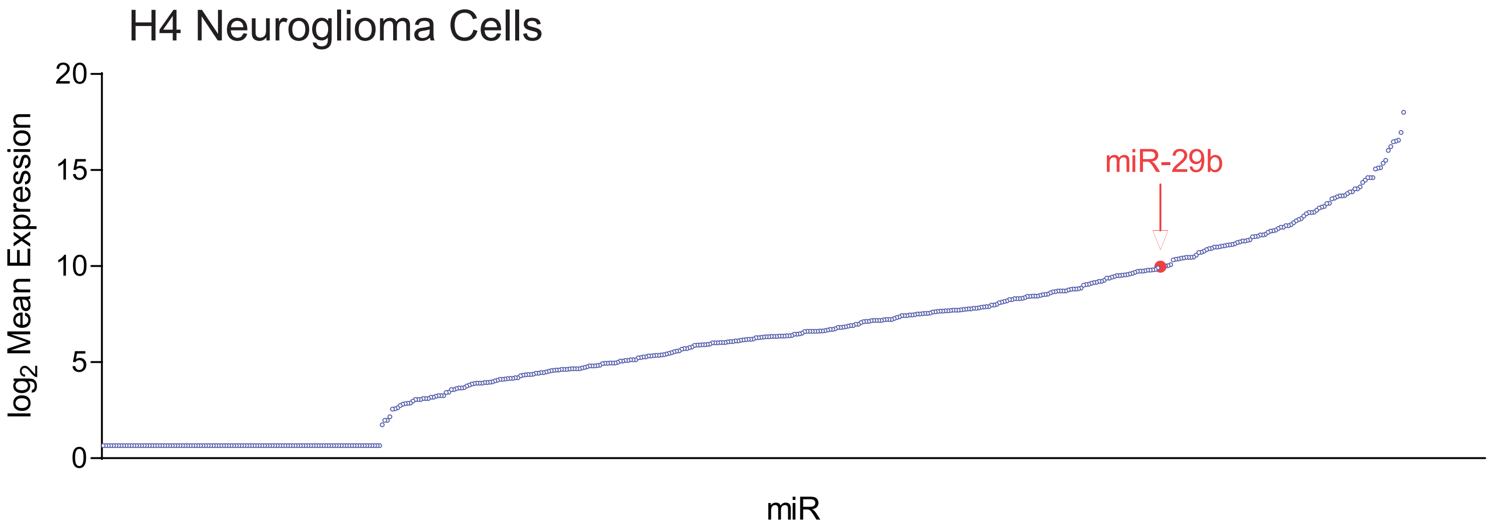

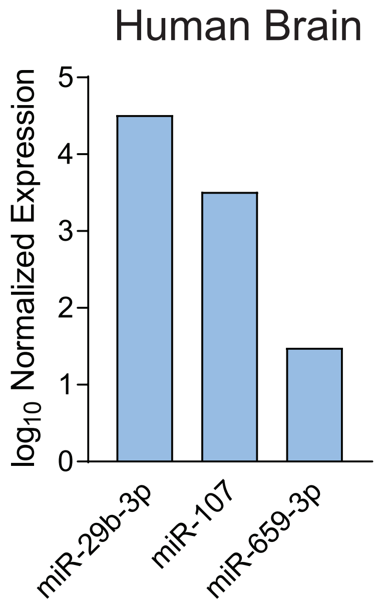

### Fig. S2

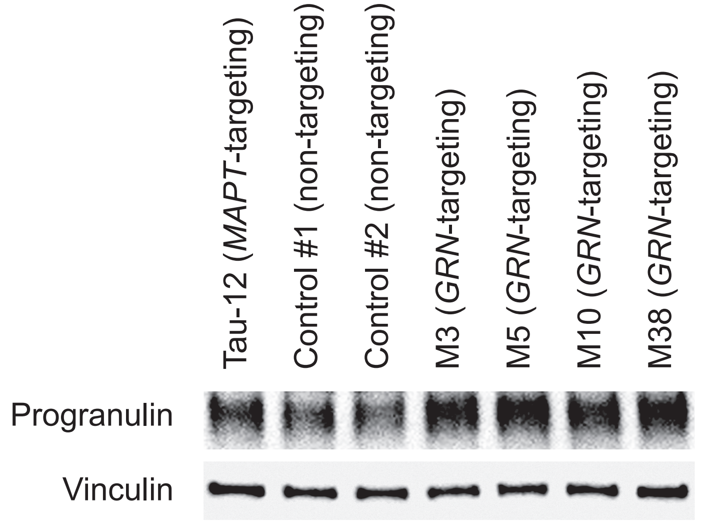

### Fig. S3

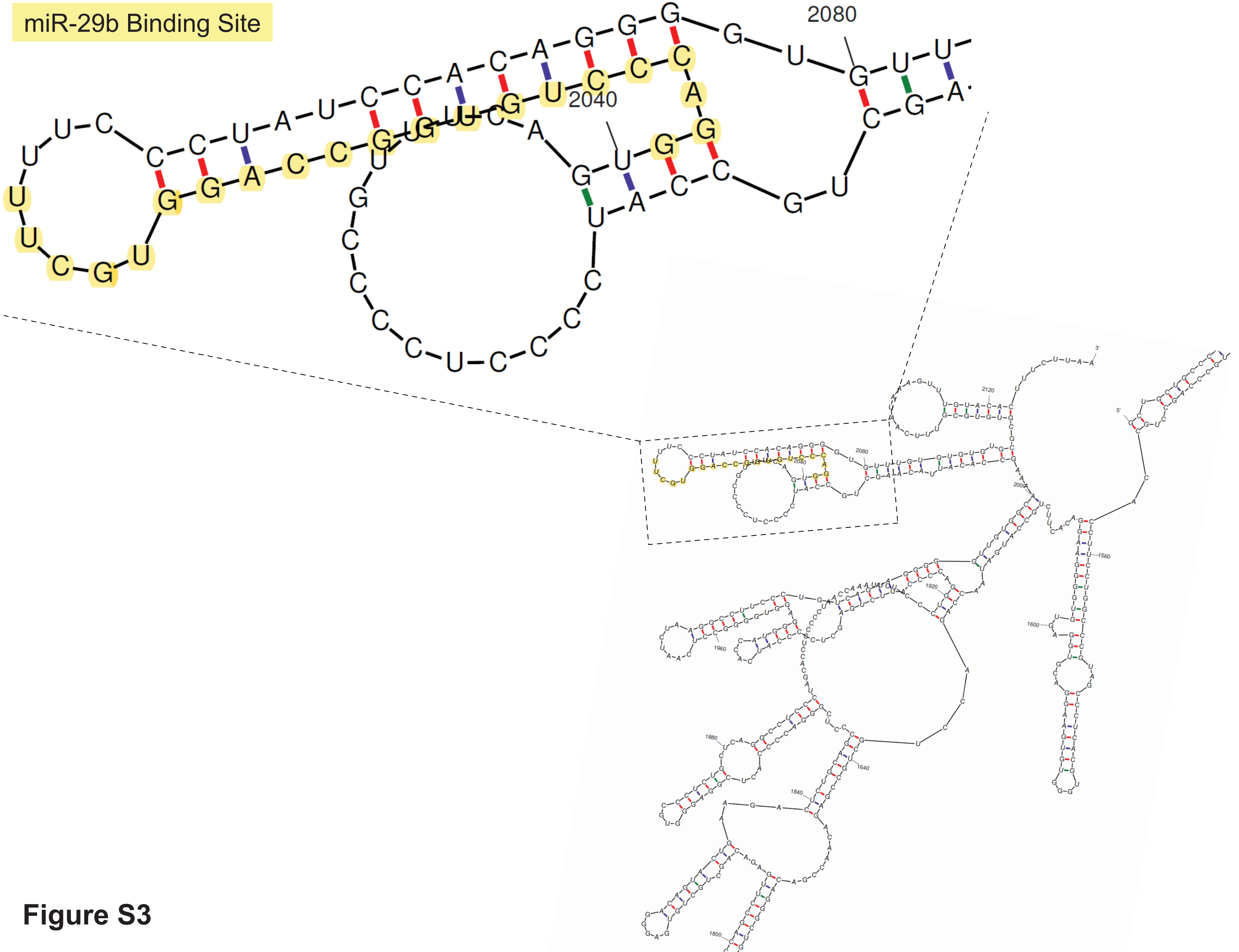

### Fig. S4

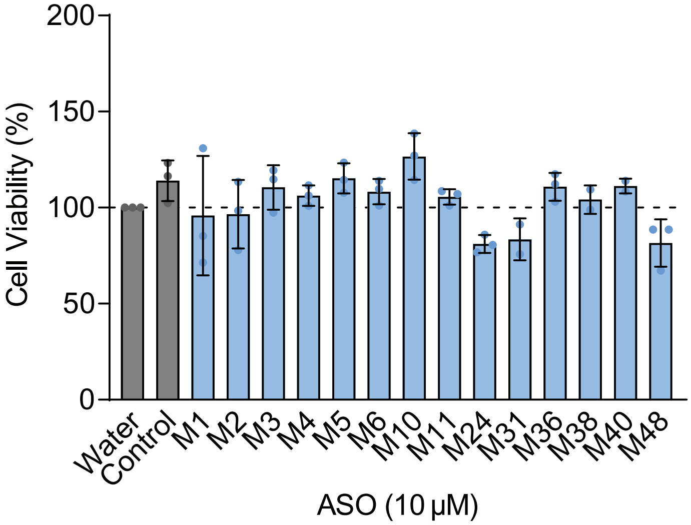

### Fig. S5

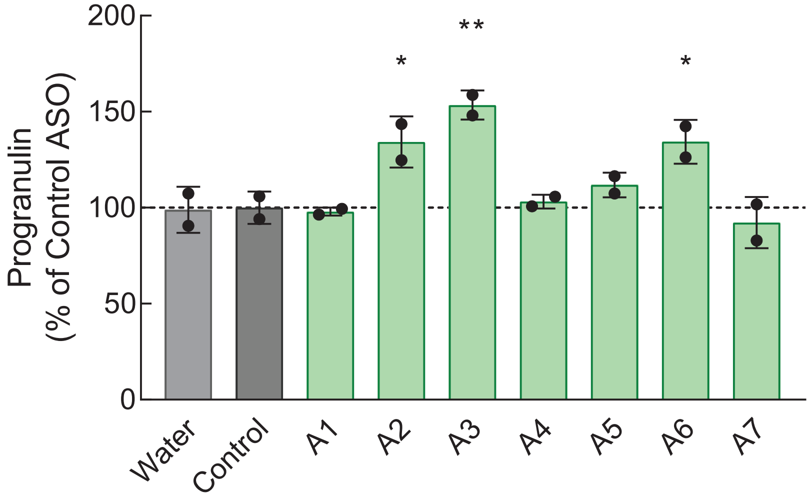

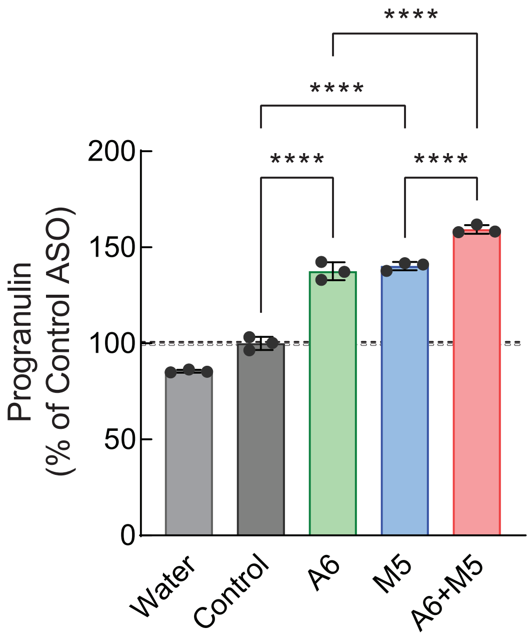
